# High pressures depress the onset of intracellular vitrification

**DOI:** 10.1101/2025.07.11.664353

**Authors:** Stewart Gault, Charles S. Cockell

## Abstract

The low temperature limit for life remains elusive and poorly understood. This ignorance is further compounded when applied to life in multi-extreme environments where low temperatures combine with factors such as high salt concentrations, or high environmental pressures. It has been proposed that the onset of intracellular vitrification enforces a biophysical low temperature limit for unicellular life at ∼ -23 ^°^C. However, it has not been demonstrated how high-pressures affect intracellular vitrification, which is vital for understanding the habitability of low temperature, subsurface environments, both on Earth and on other planetary bodies. Here, we used high-pressure differential scanning calorimetry to measure the intracellular vitrification of *Bacillus subtilis* across pressures ranging from 1 to 1000 bar. We find that high pressures depress the onset of intracellular vitrification in a pressure dependent manner, which is tightly correlated with the ability of pressure to depress the freezing point of water. Additionally, we show that sub-molar concentrations of NaCl can act in combination with high pressures to further depress intracellular vitrification, highlighting the interplay between temperature, pressure, and ions in influencing the physical state of cells in natural environments. These results show that cells in subzero high-pressure environments would be liquidous, and potentially metabolically active, and not merely vitrified and preserved. Additionally, our results provide considerations in the preparation of biological samples through high-pressure freezing for electron microscopy, particularly those associated with high concentrations of cryoprotectants.

## Introduction

The potential for active microbial activity in high-pressure subzero environments remains poorly understood. On Earth, such extreme environments are typified by Lake Vostok in Antarctica, or deep subsurface permafrost in Siberia. Lake Vostok is characterised by pressures of ∼400 bar due to 4 km of overlaying ice, and temperatures of approximately -3 ^°^C(1). In Siberia, particularly the Lena and Yana river basins, modelling suggests that the permafrost can extend to depths of ∼1.5 km, experiencing pressures over 100 bar, with environmental temperatures just below 0 ^°^C(2). The inaccessibility of these multi-extreme environments has significantly hampered our understanding of life’s ability to exist under such extremes. In contrast we know that near-surface subzero environments contain a variety of psychrophiles and cryophiles(3), with isolated species such as *Planococcus halocryophilus* remaining metabolically active at temperatures as low as -25 ^°^C(4).

In addition to the terrestrial deep subsurface, evidence suggests that planetary bodies beyond Earth harbour subzero subterranean aqueous environments. On Mars, water is either predicted to exist as deep groundwater(5), or as subglacial lakes(6,7) where temperatures would be on the order of -70 ^°^C, although the latter phenomena remains controversial. Further afield, the icy moons Europa, Enceladus, and Titan contain liquid water oceans trapped beneath their icy crusts. The icy moon oceans span a range of temperatures and pressures, with Europa and Enceladus hosting oceans at ∼0 ^°^C, but with ocean bottom pressures of 2000 bar(8) and <100 bar(9–11) respectively. Titan’s ocean however is thought to have a much broader temperature and pressure profile, with the upper water-ice boundary thought to ∼1500 bar and -14 ^°^C, whereas the ocean bottom water-rock/ice interface is thought to be ∼7500 bar and -1 ^°^C(12).

To understand the potential for such extreme environments to host life, or its remnants, we must first understand the limits to life in high-pressure, subzero environments. Attempts have been made to sample the microbial diversity of Lake Vostok, though initial measurements were significantly hampered by contamination from drill fluids(13). However, sampling of a shallower Antarctic subglacial lake, Lake Whillans, revealed a diverse microbial ecology(14–16). Ultimately though our perception of the limits to life in high pressure subzero temperature environments remains significantly limited.

In the absence of in situ measurements, the field of microbial biophysics offers a window into this otherwise challenging area of multi-extreme microbiology. From the biophysical perspective, the low temperature limit for life is thought to be enforced by the onset of intracellular vitrification, as was proposed by Clarke et al(17). Intracellular vitrification occurs when cells lose their intracellular water through osmosis as the freezing of the extracellular environment concentrates extracellular solutes. The loss of intracellular water continues until a point is reached at which intracellular diffusion ceases and a dramatic increase in viscosity occurs. The intracellular vitrification of unicellular organisms occurs at approximately -23 ^°^C in the absence of salts and other substances which can depress the freezing point of water. Cryoprotectants such as glycerol, either delivered exogenously(18) or produced in vivo(19) can significantly depress intracellular vitrification, as can the presence of environmentally relevant salts such as Mg(ClO_4_)_2_, which at 2.5 M can depress *Bacillus subtilis*’ (*B. subtilis*) intracellular vitrification to -83 ^°^C(20). While these studies show that the biophysical limits to life in subzero environments can be shifted by the environmental chemical composition, the effect of high pressures, relevant to Earth’s deep cryosphere and icy moon oceans, remains unknown.

To address this terra incognita, we have utilised high-pressure differential scanning calorimetry (HPDSC) to measure the effect of pressures up to 1 kbar on the intracellular vitrification of *B. subtilis*. These results will help elucidate the potential for life and active biochemistry in high-pressure, subzero environments and assist in the interpretation of data from missions assessing the habitability of environments beyond Earth.

## Materials and methods

*Bacillus subtilis* DSM 10 was obtained from the Leibniz Institute, DSMZ.

### Cell culturing and sample preparation

*B. subtilis* was cultured overnight in 50 mL of growth media (5 g/L peptone, 3 g/ L meat extract, pH 7, 25^°^C) in three separate flasks. The separate cell suspensions were then pelleted at 3500 g for 10 minutes, the supernatant was discarded, and the pellets were resuspended in deionized water then combined into a single cell suspension. This concentrated cell suspension was pelleted and washed twice more with deionized water. For experiments investigating the effect of NaCl on vitrification, the washed cell pellet was resuspended in the desired NaCl concentration for 1 hour before being pelleted as before. To remove excess solution from the final cell pellet, the cells were centrifuged a final time at 10,000 g for 1 minute.

### High pressure differential scanning calorimetry

High-pressure differential scanning calorimetry was conducted using a Microcalvet calorimeter with Hastelloy high-pressure cells, manufactured by Setaram with a circulating water bath to cool the HPDSC Peltier. High pressures were generated using a Teledyne LABS Syrixus 65x syringe pump with N_2_ as the pressurising gas.

For HPDSC measurements, 15 to 40 mg of *B. subtilis* cell pellets were loaded into an aluminium cup weighing 73 mg, 6 mm in height and 5 mm in diameter with walls 0.25 mm thick. The aluminium cup containing the cell pellet was placed inside the sample chamber with an identical empty aluminium cup placed inside the reference chamber. For the temperature ramps, cells were held isothermally for 5 minutes at 10 ^°^C, before being cooled to –45 ^°^C at 1 ^°^C/min, held isothermally at –45 ^°^C for 10 minutes, then heated to 10 ^°^C at 1 ^°^C/min. The HPDSC scans were conducted at 1, 250, 500, 750, and 1000 bar.

The HPDSC heating scan traces were analysed with TRIOS software (TA Instruments, Version 5.1). For analysis, the first derivative of the heating scan was produced and smoothed to 200 neighbours to highlight the intracellular vitrification signal. The vitrification signal was then analysed by recording the temperatures of the onset, peak, and end, denoted T_onset_, T_peak_, and T_end_ respectively(20).

## Results and Discussion

Here we report the first experimental investigation of the effect of high pressures on intracellular vitrification. Figure 1 shows the temperatures at which the T_onset_, T_peak_, and T_end_ of the intracellular vitrification signal was recorded across pressure. At 1 bar, *B. subtilis* cell pellets exhibited a mean T_peak_ of –24.11 ^°^C, which is consistent with previous vitrification values for *B. subtilis* and other unicellular organisms(17). As the pressure was increased, a pressure dependent depression of intracellular vitrification was observed. By 1000 bar the mean T_peak_ had been depressed to –29.78 ^°^C, with 250, 500, and 750 bar depressing T_peak_ to -25.30, -26.82, and -29.02 ^°^C respectively.

**Figure 1:**
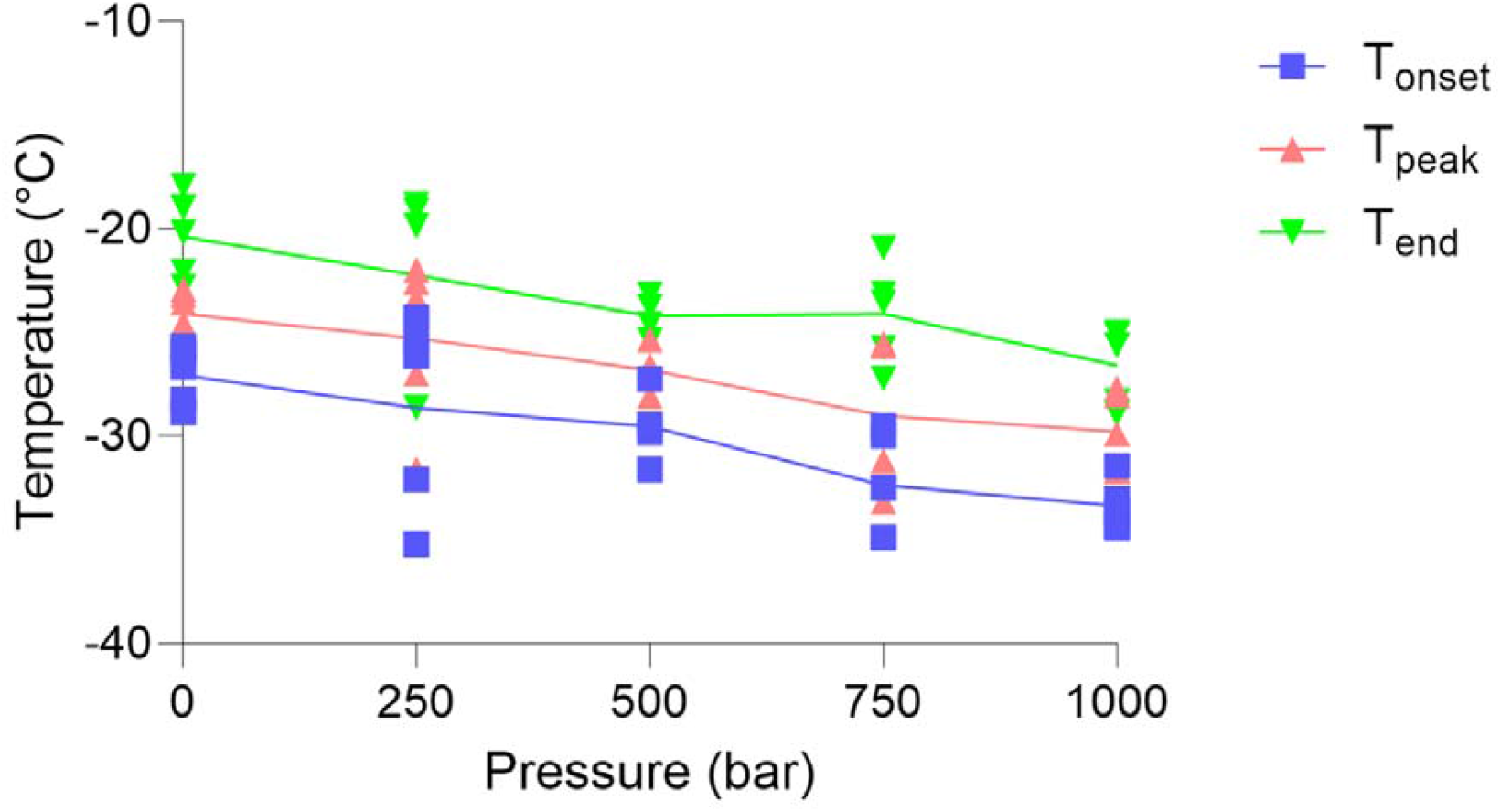
Intracellular vitrification of *Bacillus subtilis* across pressure. The T_onset_ (blue squares), T_peak_ (red triangles), and T_end_ (green triangles), of the *B. subtilis* intracellular vitrification signal at 1 bar (n = 5), 250 bar (n = 5), 500 bar (n = 4), 750 bar (n = 5), and 1000 bar (n = 5).

The gradual depression of intracellular vitrification by pressure is consistent with the pressure induced depression of water’s freezing point. This reinforces the causal relationship between the freezing of the extracellular environment and the onset of intracellular vitrification, as stipulated by Clarke et al.(17). However, the pressure induced depression of intracellular vitrification was slightly lower than what would have been expected according to the phase diagram of water. At 1000 bar, the freezing point of water is ∼-9 ^°^C, whereas intracellular vitrification was only depressed by ∼5 ^°^C.

As real environments are typically expected to contain salts, which themselves can depress the freezing point of water, we expected that the presence of salt and pressure should have a combined effect on intracellular vitrification. We demonstrate this in Figure 2 by showing that the presence of NaCl and high pressures can exhibit a combined effect on the depression of intracellular vitrification. From the representative data, at 1 bar, 0.25 M and 0.5 M NaCl depressed intracellular vitrification to ∼-24 and -32 ^°^C respectively. Increasing the pressure to 1000 bar depressed intracellular vitrification at 0 M and 0.25 M NaCl by -7 ^°^C, while a minor shift was observed in the presence of 0.5 M NaCl, with the vitrification peak being depressed by -3 ^°^C. These results suggest that the intracellular vitrification of unicellular organisms will occur at temperatures below the liquid-solid phase boundary of an environmentally relevant brine’s temperature-pressure-salt phase diagram. This is likely due to the high concentration of organic molecules within cells, and the nanoconfined nature of intracellular water(21), which may act in unison to further modulate the phase behaviour of a cell’s intracellular contents.

**Figure 2:**
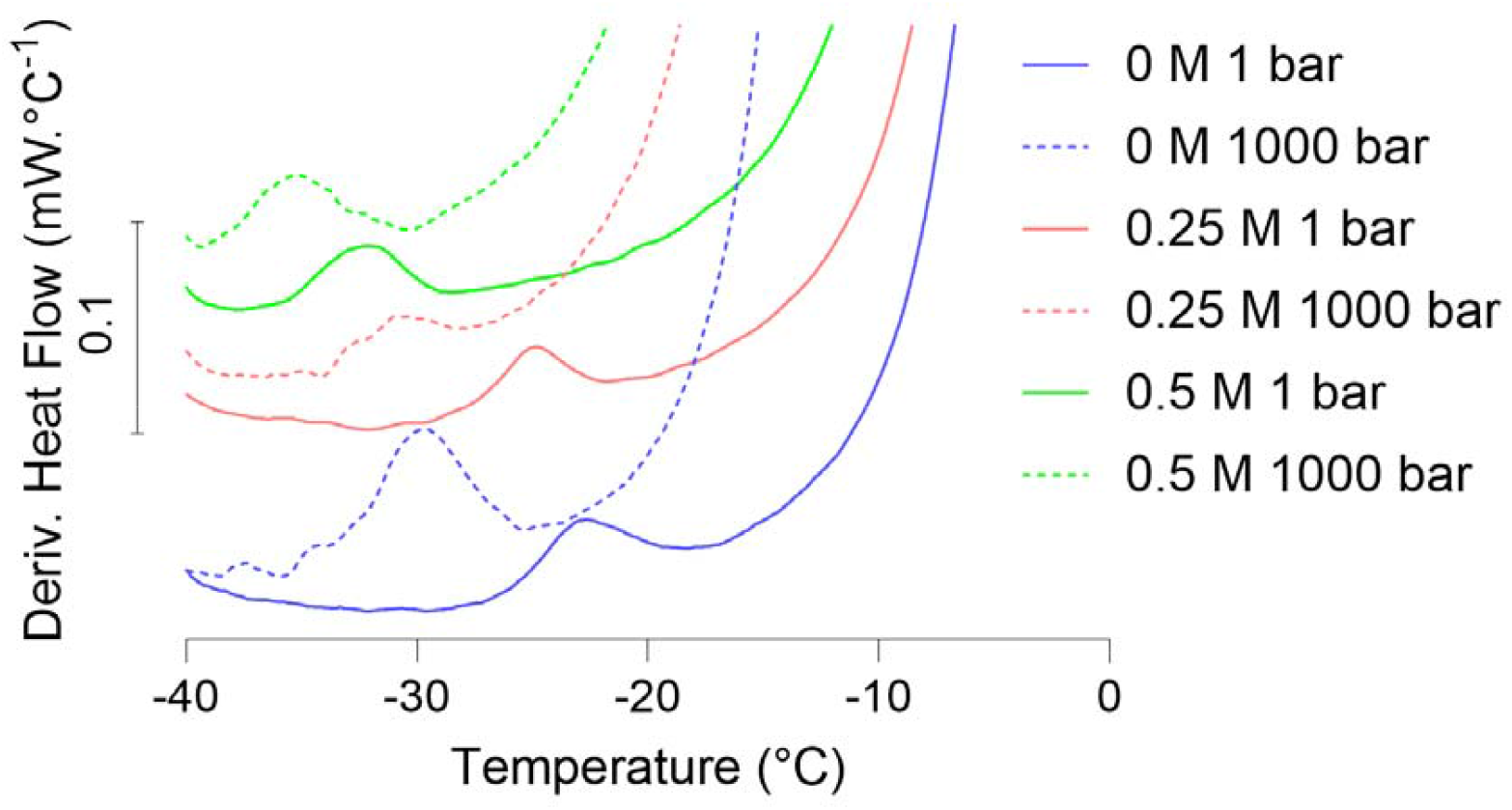
*B. subtilis* heating scans across pressure and NaCl concentrations: Representative heating scans of *B. subtilis* pellets showing the vitrification signal at 1 bar (solid lines), and 1000 bar (dashed lines) and 0 M (blue), 0.25 M (red), and 0.5 M (green) NaCl.

Due to the saturation of the heat signal caused by the mass of water in the sample required to measure vitrification, we are unable to provide a quantitative thermodynamic analysis of the melting and freezing of the water within our samples. The relatively large sample mass meant that the freezing/melting signal of water was consistently saturating the calorimeter, however qualitatively it was observed that the freezing and melting of water was depressed by high pressures and salt concentrations, as would be expected with their phase diagrams.

These results suggest that unicellular organisms in subzero high-pressure brines would remain in a fluid state under environmental conditions analogous to those experienced in the ice surrounding Lake Vostok, deep subsurface permafrost, and the ice shells of Europa and Enceladus, close to the ocean-ice boundary. However, our conceptual understanding of whether an environment is habitable or not is extremely biased by our understanding of the biophysical and biochemical limits to terrestrial life. So, while Lake Vostok is habitable from our theoretical understanding of life’s biophysical limits, it is yet to be demonstrated whether microbial life itself can remain active in these cold, high-pressure, nutrient poor environments. In the context of high pressure, subzero aqueous environments like those of Mars, Europa, Enceladus, and Titan, the lack of representative, accessible terrestrial analogues to these multi-extreme environments further compounds our ignorance as to their habitability.

As the microcalorimeter’s highest operating pressure is 1 kbar we were unable explore the pressures that are expected at the base of Europa and Titan’s oceans, 2 kbar and 7.5 kbar respectively(8,12). The potential for brine veins in high-pressure ice to support life is further tempered by our understanding of relationship between pressure and biochemical structure. Pressures of 7.5 kbar are sufficient to denature proteins(22,23), as such, life in these environments would necessitate biochemical adaptations and innovations that life on Earth has not experienced in its evolutionary history. The lower operational temperature of -45 ^°^C also precluded examining environmental conditions analogous to the Martain subsurface where pressures are expected to be <1 kbar, but the temperature can be as low as -70 ^°^C. Such data would have complimented previous work where we showed that molar concentrations of Mg(ClO_4_)_2_ can significantly depress *B. subtilis*’ vitrification, with 2.5M Mg(ClO_4_)_2_ depressing T_peak_ to ∼-83 ^°^C. Given that intracellular vitrification is closely linked to the freezing points of aqueous solutions, fully mapping the high-pressure phase diagrams of environmentally relevant ionic solutions will greatly expand our understanding of where the biophysical low temperature limits for life may be found in extreme environments in our solar system.

Our results may also impact protocol design for electron microscopy sample preparation. High-pressure vitrification and freezing is a mainstay for the cryofixation of electron microscopy samples, wherein small volumes of samples are sealed then rapidly cooled(24). As the water within the sample begins to freeze, the ensuing volumetric expansion causes a rapid increase in the internal pressure of the sample which inhibits ice crystallisation, instead inducing the formation of amorphously frozen/ vitrified water, allowing for greater preservation of biological structures. Until now however it had not been demonstrated what effect high pressures have on cellular vitrification. Therefore our results suggest that the process of high-pressure freezing will not only depress ice crystal formation, but may also affect the temperature at which the cells themselves undergo a glass transition. Given the demonstrated relationship between pressure and ion composition on intracellular vitrification, it stands that high concentrations of deeply eutectic ions may affect the degree to which samples can fully vitrify, thus affecting sample quality for electron microscopy. This would likely affect the sample preparation of halophiles which require high salt concentrations for optimal growth, or species which can produce high concentrations of cryoprotectants in vivo(19), which may need to be brought to lower temperatures to achieve a sufficient degree of vitrification.

## Acknowledgements

I would like to thank Dr Claire Hobday and Dr Joshua Levinsky for providing access to and training for the high-pressure differential scanning calorimeter and Toby Fu for machining the aluminium cup.

## Data Availability

All data for Figures 1 and 2 are provided in a supplementary file.

